# Reduced skeletal density and mechanical resistance in mangrove-associated *Pocillopora* sp. corals

**DOI:** 10.64898/2026.10.07.757461

**Authors:** Adi Zweifler (Zvifler), Dayana Chadda-Harmer, Matthew Foley, Shawna A. Foo

**Author notes:** Corresponding author, Adi Zweifler (Zvifler).

## Abstract

Coral skeletal integrity underpins reef habitat complexity and coastal protection, yet it remains unclear whether corals persisting in chemically variable marginal habitats, such as mangrove-associated reefs, maintain sufficient mechanical resistance to withstand physical disturbance. We combined micro-computed tomography (micro-CT) morphometric analysis with compression testing to quantify skeletal microarchitecture and compressive mechanical performance of *Pocillopora* sp. skeletons collected across a habitat gradient at Low Isles, Great Barrier Reef, from mangrove-associated sites to exposed reef sites. Baseline skeletal properties varied among sites, with the Proximal Reef site having the densest and least porous skeletons, while mangrove-associated sites had the most porous skeletons. Similarly, Proximal Reef skeletons exhibited two-to-three-fold greater compressive stiffness and peak force than mangrove-associated skeletons. Percentage object volume was a stronger predictor of compressive stiffness than surface area-to-volume (S/V) ratio, indicating that bulk skeletal density better captured load-bearing capacity than surface architectural complexity. Structural failure mode also differed markedly among habitats: mangrove-associated skeletons underwent catastrophic fragmentation of their open, porous framework, whereas Distal and Proximal Reef skeletons failed through discrete crack-plane propagation while largely maintaining overall structural coherence. These results indicate that the physiological stress tolerance associated with mangrove habitats occurs alongside reduced mechanical resistance and more destructive failure modes, suggesting a potential trade-off between chemical stress tolerance and structural investment. Assessments of resilience in marginal coral populations should therefore consider skeletal mechanical performance alongside physiological tolerance, particularly as climate change intensifies storms and alters hydrodynamic disturbance regimes.

## Introduction

Coral skeletal integrity underpins the primary functional roles of reef ecosystems, including structural habitat complexity, coastal protection, and the three-dimensional framework on which reef biodiversity depends (Graham and Nash, 2012). The persistence of these ecosystem functions depends on the ability of coral colonies to maintain structural stability over time, resisting bioerosion, supporting reef accretion, and providing habitat for a diverse range of reef-associated organisms (Graham and Nash, 2012; Perry et al., 2018). Consequently, variation in skeletal structure and strength can have important implications not only for individual colony survival but also for the long-term functioning and resilience of coral reef ecosystems (Alvarez-Filip et al., 2009; Omer et al., 2020).

As climate change accelerates, coral reefs are increasingly exposed to both chronic stressors, such as ocean warming and acidification (Anthony, 2016; Hoegh-Guldberg et al., 2007), and acute disturbances, including more intense tropical cyclones and storm-driven wave action (IPCC, 2023). These stressors can alter coral calcification, skeletal density, and microstructural organisation, potentially reducing the capacity of colonies to withstand mechanical loading and physical damage (Madin et al., 2014; Mollica et al., 2018). Resistance to such disturbances is governed by both material properties (e.g. density, porosity), and architectural characteristics (e.g. skeletal microstructure, colony morphology) (Moynihan et al., 2021). Understanding how environmental conditions shape these traits is therefore critical for predicting reef persistence under future climate scenarios.

Coral skeletons are formed by the deposition of calcium carbonate (aragonite) and organic matrix molecules by calicoblastic cells located on the aboral ectoderm of coral polyps (Capasso et al., 2022; Milita et al., 2023). Aragonite is initially precipitated as granular crystals from which elongate, needle-like crystals develop to form a sclerodermite, the fundamental building block of coral skeletons (Cohen and Holcomb, 2009; Coronado et al., 2019). Sclerodermites are subsequently accreted into elongate chains that form micro- and macroscopic skeletal structures such as pores and channels within the skeleton (Al-Horani, 2015).

Coral skeletal strength is influenced by a range of structural and compositional properties, each describing a specific aspect of mechanical performance and captured using different measurement approaches. Bulk density, typically measured via buoyant weight (Davies, 1989) or micro-computed tomography (micro-CT) derived percentage object volume (Bertini et al., 2026), reflects the overall proportion of mineralised material within a given skeletal volume. Porosity describes the complementary proportion of non-mineralised space and is quantified from micro-CT reconstructions alongside surface-area to volume (S/V) ratio, structure thickness, and structure separation (Chadda-Harmer et al., 2025a). Together, these parameters describe the geometry and connectivity of the skeletal framework. Beyond architecture, skeletal composition can be characterised through organic matrix content, reflecting the proportion of organic material embedded within the aragonite mineral phase, and through aragonite crystal orientation, measured using techniques such as electron backscatter diffraction or crystal orientation mapping, which capture the crystallographic alignment of the skeleton’s mineral building blocks (Moynihan et al., 2022). Each of these properties contributes to mechanical performance by influencing either the amount of load-bearing material present (bulk density, porosity), the thickness and connectivity of load-bearing elements (S/V ratio, structure thickness), or the intrinsic material properties of the skeleton itself (organic matrix content, crystal orientation). For example, higher organic matrix content has been associated with increased skeletal fragility (Moynihan et al., 2021).

Despite reflecting fundamentally different aspects of skeletal organisation, these properties are often measured in isolation. As a result, microstructural metrics such as porosity, S/V ratio, and structure thickness are frequently used as proxies for skeletal strength, with more porous or higher S/V skeletons generally assumed to provide lower mechanical strength. However, whether these microstructural differences reliably translate into differences in actual compressive performance remains poorly resolved (Moynihan et al., 2021, 2022; Gibson & Ashby, 2010; Keaveny et al., 2001). These properties can also vary independently of one another and do not always align with intuitive assumptions about strength. For example, a coral skeleton may exhibit high S/V due to many fine pores while retaining thick skeletal struts and high mineral density, thereby maintaining substantial compressive strength (Madin et al., 2014). Conversely, low S/V may mask extensive internal void space or thin load-bearing walls that reduce structural integrity. This decoupling between microstructural architecture and mechanical performance highlights a critical limitation of relying on a single architectural metric to infer compressive strength (Figure 1) (Moynihan et al., 2021; Pasquini et al., 2015).

**Figure 1.**
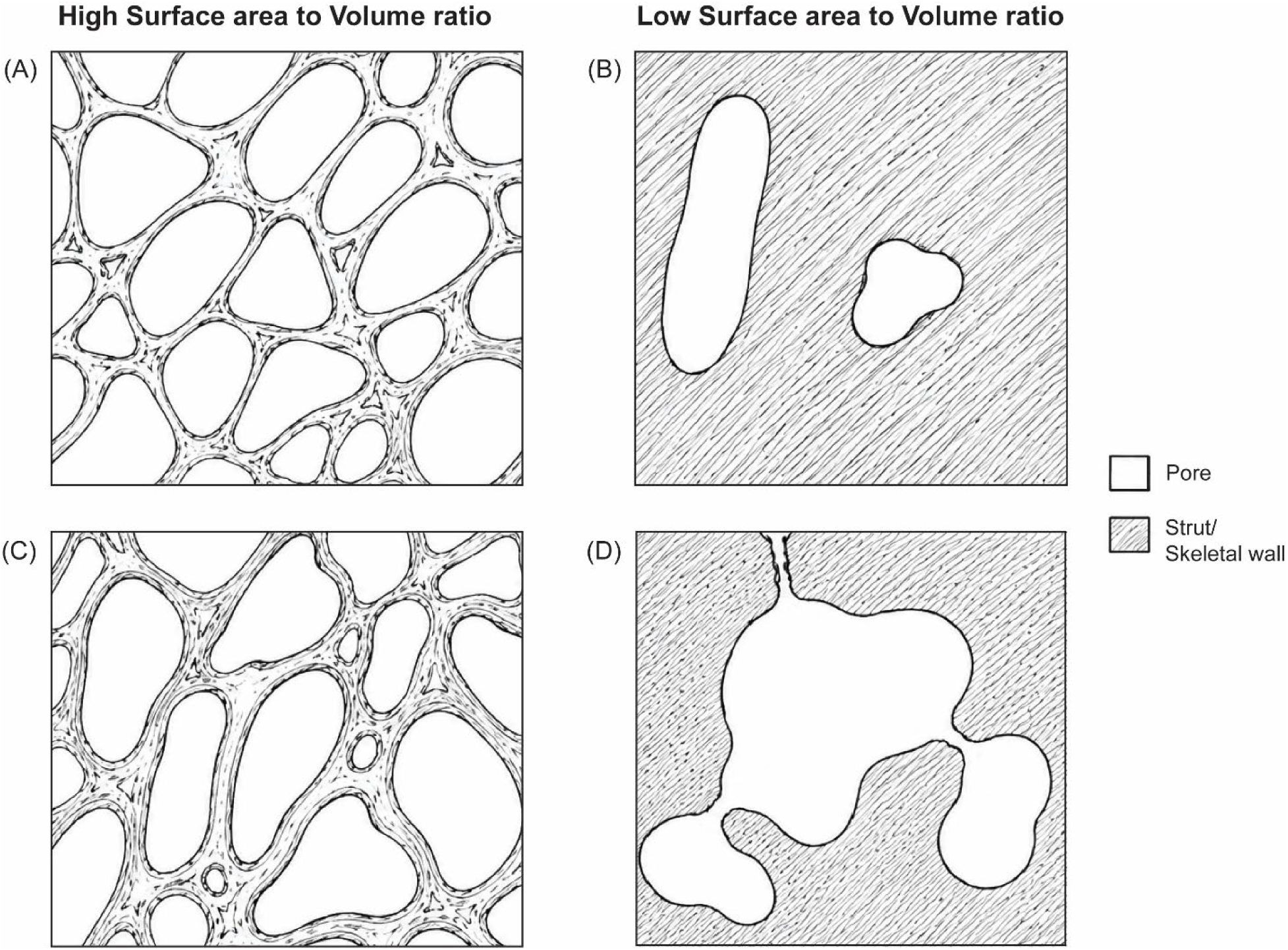
Conceptual illustration of the potential decoupling between surface area-to-volume (S/V) ratio and compressive strength in coral skeletal architecture. Panels contrast high S/V (A, C) and low S/V (B, D) structural configurations. (A) High S/V architecture characterised by numerous pores separated by relatively thin skeletal walls, resulting in extensive internal surface area and less load-bearing material. (B) Low S/V architecture with simple, isolated pores separated by thick skeletal walls, representing the conventional expectation that a lower S/V ratio reflects a denser and potentially stronger structure. (C) An alternative high S/V architecture in which numerous pores coexist with relatively thick skeletal walls, demonstrating that high S/V does not necessarily imply low structural strength. (D) An alternative low S/V architecture in which a large, interconnected void system is surrounded by predominantly thick skeletal material but includes localised thin walls or narrow connections, creating potential points of structural weakness. Together, panels C and D illustrate how a single average S/V value can conceal important variation in internal architecture and therefore may not reliably predict compressive performance. White regions denote pore space; hatched regions denote mineralised skeletal material (strut/skeletal wall).

Hydrodynamic regime is a major environmental driver of coral skeletal morphology and mechanical performance (Madin et al., 2008; Madin and Connolly, 2006). Coral inhabiting high-energy environments are frequently exposed to strong wave action and hydrodynamic loading, which can favour morphologies with denser skeletons that enhance resistance to breakage and dislodgment (Madin et al., 2014; Madin and Connolly, 2006). Water motion may also enhance calcification by increasing mass transfer at the coral-water interface, promoting nutrient delivery and the removal of dissolved CO₂ and metabolic waste products from the coral tissues (Comeau et al., 2014; Falter et al., 2013; Nakamura and Van Woesik, 2001). In contrast, corals in hydrodynamically protected environments, such as mangrove lagoons, likely experience reduced selection for maximum skeletal strength, potentially favouring investment in traits that enhance tolerance to thermal, chemical, and physiological stress, including increased physiological plasticity and metabolic flexibility (Camp et al., 2019; Scucchia et al., 2023). However, the mechanical consequences of this trade-off for physical disturbance resistance remain poorly characterised. Other abiotic stressors associated with climate change, particularly ocean acidification, are also known to alter skeletal microstructure and crystal organisation. Lower pH conditions have been linked to increased skeletal porosity, thinner skeletal elements and altered aragonite crystal organisation in corals (Coronado et al., 2019; Tambutté et al., 2015).

Despite widespread reef degradation, some coral populations persist in naturally extreme environments that experience conditions resembling those projected for future oceans (Haydon et al., 2023; Scucchia et al., 2023). Mangrove lagoons are often characterised by elevated temperatures, high pCO₂ concentrations, reduced pH, and highly variable carbonate chemistry (Camp et al., 2017; Waller et al., 2025; Yates et al., 2014). The continued persistence of corals in these habitats suggests that long-term exposure to environmental extremes may promote physiological acclimatisation, local adaptation, or the selection of skeletal traits that enhance survival under stress (Camp et al., 2019, 2018). As such, corals from naturally extreme or marginal environments provide a valuable opportunity to investigate how chronic environmental stress influences skeletal architectures and mechanical performance, offering valuable insights into the mechanisms underpinning coral resilience in a changing climate. Evidence suggests that these environments can substantially alter coral skeletal structure. For example, *Porites lutea* colonies growing in mangrove lagoons exhibit reduced skeletal density and thickness alongside increased porosity relative to adjacent reef habitats (Scucchia et al., 2023). Similarly, corals from naturally elevated CO_2_ vent sites show increased skeletal porosity together with reduced skeletal density and stiffness (Fantazzini et al., 2015; Prada et al., 2017). Although these structural changes are widely interpreted as increasing susceptibility to mechanical damage, this conclusion has generally been inferred from skeletal density and porosity rather than evaluated through direct mechanical testing. Consequently, it remains unclear whether the microstructural modifications observed in naturally extreme environments translate into reduced compressive strength, or whether compensatory traits, such as greater skeletal connectivity or increased organic matrix content, mitigate these potential trade-offs. Resolving this uncertainty is essential for predicting which coral populations are most likely to withstand both chronic environmental stress and acute physical disturbances.

Prior attempts to characterise coral skeletal mechanical properties have largely relied on destructive compression testing of intact fragments without accompanying microstructural analyses. Other methods such as nanoindentation and other microscale techniques, quantify local material properties (e.g., hardness and elastic modulus) at the scale of individual skeletal elements rather than whole-fragment mechanical behaviour (Pasquini et al., 2015). More recently, micro-CT has been combined with compression testing to directly link three-dimensional skeletal architecture with whole-fragment mechanical performance. However, this approach has only been demonstrated on a single coral fragment and has not yet been applied to compare skeletal mechanical performance across environmental gradients (Altintas et al., 2024).

In this study we combined micro-CT imaging with compression testing to quantify the compressive mechanical properties of *Pocillopora* sp. skeletons collected across a mangrove to reef gradient at Low Isles, Great Barrier Reef. Specifically, we aimed to (1) quantify compressive stiffness (the resistance to elastic deformation under load) and peak force (the maximum load sustained before structural failure) of coral skeletons across four reef sites spanning a gradient from mangrove lagoon to reef flat, (2) determine whether habitat-driven differences in skeletal microarchitecture translate to measurable differences in mechanical properties, (3) assess the relationship between baseline skeletal porosity and compressive stiffness, explicitly comparing S/V ratio and percentage object volume as complementary proxies of porosity and (4) characterise the failure modes and microstructural deformation patterns associated with compression, and how these differ among habitat types. By integrating stepwise micro-CT scanning with mechanical loading of the same samples, this approach enables direct quantification of structural changes alongside force-displacement behaviour. While a similar method has been previously applied to bulk coral reef limestone (Altintas et al., 2024; Meng et al., 2022), to our knowledge, this is among the first applications of this combined approach to individual coral skeleton fragments, and the first to compare mechanical performance of coral fragments collected across an environmental gradient. Given trade-offs between stress tolerance and structural investment, we hypothesised that coral skeletons closer to, or within, the mangrove system would be mechanically weaker than those from the adjacent open reefs, reflecting reduced selection for skeletal robustness under lower energy, but chemically stressful conditions.

## Materials and Methods

### Study site and sample collection

*Pocillopora* sp. (Lamarck, 1816) coral samples were collected from four reef sites at Low Isles, a coral cay and mangrove island approximately 15 km north-east of Port Douglas, Queensland, Australia (Figure 2). The four sites span a gradient from enclosed mangrove habitat to reef flat: (1) Mangrove Lagoon (n = 4), (2) Mangrove Channel (n = 4), (3) Proximal Reef (n = 4), and (4) Distal Reef (n = 3). Mangrove Lagoon and Mangrove Channel are located within the protected mangrove system, characterised by low oxygen and energy conditions and elevated temperatures and *p*CO₂ relative to the open reef (Haydon et al., 2021). The Mangrove Channel site experiences tidal flow, consistent with the atypical, mud-poor sediment grain size assemblage previously described for tidally influenced areas of Low Isles (Frank and Jell, 2006; Hamylton et al., 2019). Proximal and Distal Reef sites (in relation to the mangrove-associated sites) are located on the inner and outer reef margin, respectively, representing more exposed reef environments. Sample sizes reflect the availability of suitable samples, as well as the destructive and time intensive nature of compression testing.

**Figure 2.**
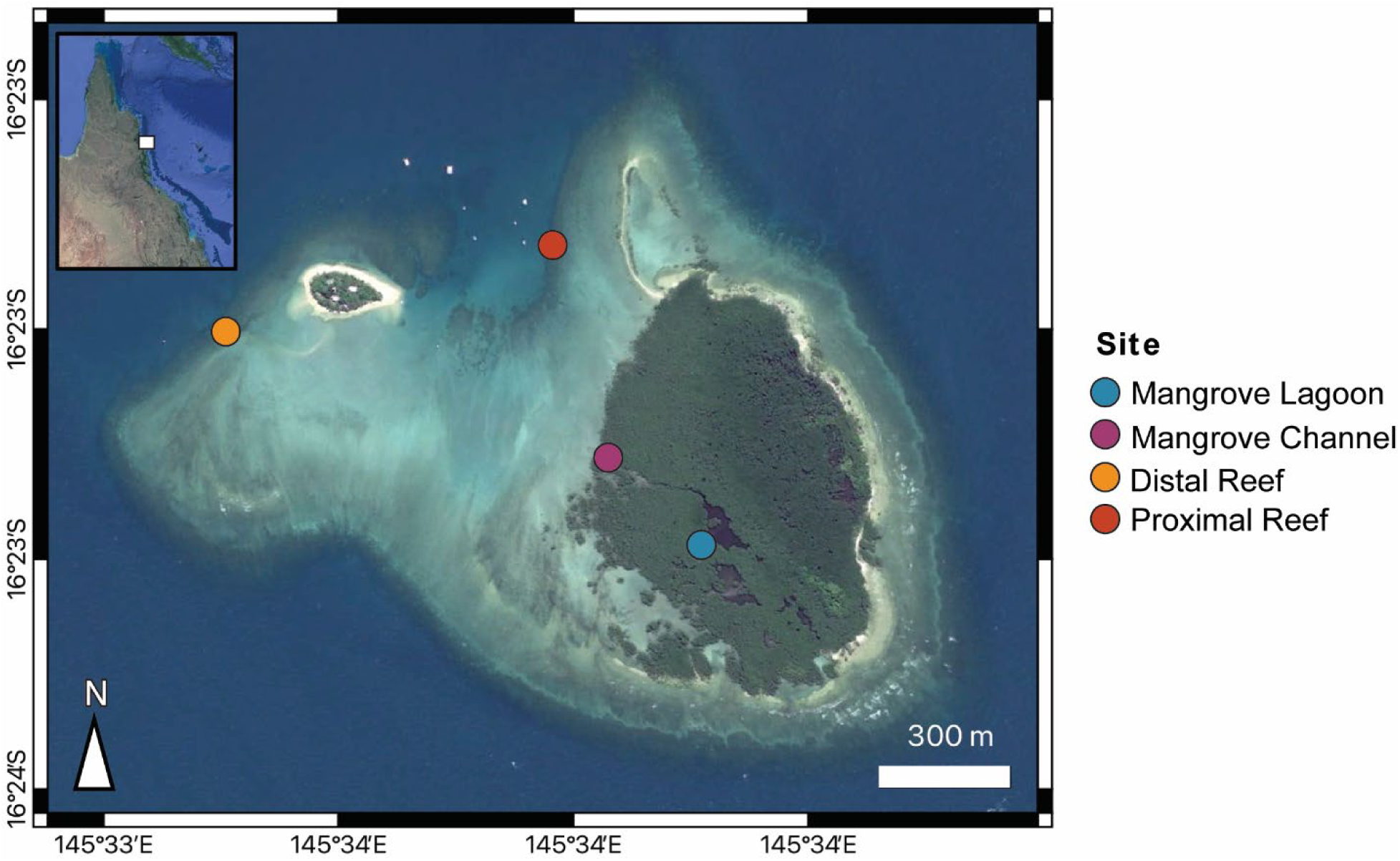
Location of Low Isles on the Great Barrier Reef, Australia including map of sampled sites (Google Earth).

### Sample preparation and compression testing

Coral fragments were sectioned using a hand-held mini rotary tool, cutting a 1 cm piece from the branch tip to ensure consistent sections across all samples. Opposing faces were polished to ensure flat, parallel surfaces for uniform load distribution during compression. Compression was applied axially along the branch’s long (growth) axis, with force applied vertically downward from the top of the sample onto the upper polished face (Altintas et al., 2024). The same standardised ∼1 cm section length and loading orientation were used for all samples.

Compression testing was performed using a Bruker SkyScan 1272 micro-CT system (Bruker MicroCT, Kontich, Belgium) equipped with an integrated material testing stage (MTS; maximum capacity 4400 N). The compression stage was mounted directly on the system’s rotation stage and controlled externally to the system. Two loading approaches were evaluated: (i) force-controlled loading with incremental steps, and (ii) displacement-controlled loading at a constant rate. Force-controlled loading was used for three samples (Table S1). Peak force and stiffness values from these samples fell within the range observed for displacement-controlled from the same site, confirming that the loading method did not introduce bias in the mechanical measurements. Displacement-controlled loading was adopted as the primary protocol for the remaining samples because it allowed specimens to fail progressively rather than catastrophically, while maintaining consistent deformation rates across samples of varying stiffness. At predefined displacement intervals, loading was paused to acquire static tomographic scans, preventing motion artefacts. Projections were acquired at 80kV with a Al 0.5 mm + Cu 0.038 mm filter at an isotropic voxel of 20 μm across all loading steps. Three-dimensional volumes were reconstructed using Bruker NRecon software. Scans acquired at each pause during compression testing were subsequently used for the morphometric analysis described below.

### Micro-CT morphometric analysis

Three scans were taken per sample: a pre-compression baseline (T1), an intermediate scan taken near but before peak loading (e.g., failure) (T2), and a final scan at or after failure (T3), yielding 45 scans in total (n=15 samples X 3 scans per sample). The T2 scan does not represent an equivalent loading state across samples. Because the exact force or displacement required for failure could not be known in advance, each sample was scanned as close to failure as possible. Consequently, the scans correspond to varying compressive loads and degrees of deformation based on each sample’s unique mechanical response. The fraction of total loading elapsed by T2 ranged from approximately 67% to 95%. Therefore, T1 vs T3 comparisons were the primary focus of the morphometric analysis.

Three-dimensional morphometric parameters were extracted from each scan using CT-Analyser (CTAn) (Bruker MicroCT) within a fixed volumetric region of interest, which was consistent across samples. Eleven parameters spanning volumetric, surface-based, and strut-level measures of skeletal architecture were quantified (Table 1). Absolute and percentage changes were computed between T1 and T3 for each parameter. The change between T1 and T3 quantifies the extent of compression-induced structural degradation within each sample, independent of its baseline architecture, and is distinct from the between-site baseline comparisons described below. Since distinguishing between surface-based and volume-based measures of porosity is essential for interpreting structure-property relationships in coral skeletons, in this study we explicitly compare S/V with percentage object volume as complementary proxies of skeletal porosity to evaluate how different aspects of microstructure contribute to compressive stiffness and failure behaviour. The S/V ratio captures surface-connected architectural complexity, whereas percentage object volume reflects bulk skeletal mineral content (i.e., volume fraction), which is a primary determinant of mechanical stiffness and strength in cellular and skeletal materials (Gibson, 2003; Keaveny et al., 2001). In contrast, structural-based metrics do not directly quantify load-bearing material content and therefore may not reliably predict compressive mechanical performance.

**Table 1.** Morphometric parameters and their interpretive relevance to coral skeletal properties.

| Parameter | Unit | Description | Biological/mechanical interpretation |
| --- | --- | --- | --- |
| Object volume | mm<sup>3</sup> | Total volume of skeletal (solid) material | Absolute amount of mineralised material deposited; reflects calcification extent within the sampled region |
| Percentage object volume | % | Fraction of the sample occupied by skeletal material (bulk volume fraction) | Primary proxy for bulk mineral density/porosity; strongly predicts compressive stiffness and strength in cellular materials |
| Object surface area | mm<sup>2</sup> | Total surface area of the skeletal structure | Reflects architectural complexity and surface exposure; increases with microcracking or fragmentation |
| Intersection surface area | mm<sup>2</sup> | Surface area at junctions/connections between skeletal elements | Indicates connectivity of the load-bearing network; declines with disconnection or fracture propagation |
| Surface to volume ratio | mm<sup>-1</sup> | Object surface area relative to object volume | Proxy for surface-connected architectural complexity/porosity; higher values indicate finer, more porous struts rather than bulk density |
| Object surface density | mm<sup>2</sup>/mm<sup>3</sup> | Surface area normalised to the total sample volume (rather than object volume)<sup>1</sup> | Complements S/V ratio; sensitive to how much surface is packed into a given tissue volume, independent of solid fraction |
| Structure thickness | mm | Mean thickness of skeletal struts | Indicates robustness of individual load-bearing elements; thinning suggests crushing or dissolution |
| Structure separation | mm | Mean distance between skeletal structures (pore/void size) | Proxy for inter-skeletal pore space; decreases as elements compact together under load |
| Structure linear density | mm<sup>-1</sup> | Number of structure/void transitions per unit length | Reflects how finely subdivided the skeletal network is; increases with fragmentation |
| SD of structure thickness | mm | Variability in strut thickness across the sample | Indicates heterogeneity of the load-bearing network; stable values suggest uniform damage rather than localised weakening |
| SD of structure separation | mm | Variability in pore/void size across the sample | Indicates spatial heterogeneity of porosity; stable values suggest compression alters the mean structure without changing its spatial pattern |

<sup>1</sup> Total sample volume represents the entire volume within the region of interest, including both mineralised skeletal material and pore space, whereas object volume represents only the portion occupied by mineralised skeletal material.

### Stiffness calculation

Force and displacement were recorded continuously throughout compression by the MTS load cell and displacement sensor. The intermittent micro-CT scan pauses used to acquire T1 and T3 imaging did not interrupt this mechanical record. These curves show the applied compressive force on each coral sample and resulting displacement during loading. Structural stiffness was determined from the slope of the linear elastic region of each force-displacement curve. The initial linear region represents elastic deformation, where the skeleton deforms proportionally to the applied load and would recover if unloaded. This is followed by a peak and drop in force indicating structural failure. In practice, we identified failure as the point where each sample first showed a sudden and sustained loss of load-bearing capacity, distinguishing true skeletal failure from the temporary unloading used during the scanning protocol.

The analysis was conducted in two stages: first, identifying the loading phase, defined as the portion of the continuous force-displacement record preceding the first genuine structural failure. Second, extracting the linear elastic region within this phase, to calculate stiffness (described further below). Since our samples span a wide range of applied forces (237 N to ≥3331 N), a fixed absolute force threshold was inappropriate for identifying failure. The stepwise loading protocol also included brief force reductions during pauses for micro-CT imaging, which occurred repeatedly throughout the test at predefined displacement intervals and needed to be distinguished from genuine fracture events. These planned unloading periods represented only 2–15% of the concurrent applied force, whereas structural failure produced decreases of ≥20%. Therefore, unloading was assessed relative to the applied force rather than using an absolute threshold. The loading phase was terminated at the first data-point interval satisfying both: (i) an absolute force decrease of ≥50 N *and* (ii) a relative decrease of ≥20% of the concurrent force. This combined criterion correctly distinguished all planned unloading periods from structural failure events across all 15 samples. Loading was not continued beyond the first genuine structural failure event.

The ‘linear elastic region’ was extracted as data points with force between 20% and 70% of the loading-phase maximum. The 20% lower bound excludes the non-linear region caused by initial seating of the sample against the compression plates and initial crack closure. The 70% upper bound excludes the yield zone near peak load. An ordinary least-squares regression of force on displacement was fitted to this window with the slope taken as stiffness (N mm⁻¹). The coefficient of determination (R²) was recorded and samples with R² < 0.98 were flagged as having a potentially non-linear elastic region (Table S1).

Stiffness was correlated with T1 values of all eleven morphometric parameters listed in Table 1 using Pearson’s *r* and Spearman’s ρ. Both parametric and rank-order measures were computed because Proximal Reef samples occupied extreme positions for several of these parameters. This allowed us to check whether the correlation reflected a trend across all sites or was driven primarily by the separation between Proximal Reef cluster and the rest of the samples.

### Statistical analysis

Changes in morphometric parameters from T1 to T3 were assessed using Wilcoxon signed-rank tests pooled across all samples irrespective of site (n = 15 pairs). Differences in baseline (T1) morphometric parameters among sites were assessed with the Kruskal-Wallis test (n = 15). Pairwise post-hoc comparisons used Dunn’s test (Bonferroni correction) for morphometric parameters and Mann-Whitney U tests (Bonferroni correction, six comparisons) for skeletal porosity. Between-site stiffness differences were also evaluated with the Kruskal-Wallis test. Correlations between stiffness and baseline morphometric parameters were assessed using Pearson’s r and Spearman’s ρ, with resulting p-values adjusted for multiple comparisons across the eleven parameters using the Benjamini-Hochberg procedure. All statistical analyses were conducted in R (v4.4.2) using dplyr, ggplot2, dunn.test, pheatmap, and ggbeeswarm. Statistical significance was set at α = 0.05.

## Results

### Baseline skeletal microstructure differs across sites

Certain baseline skeletal microstructure parameters differed significantly among sites (Table S2 and S3). Kruskal-Wallis tests indicated significant differences in the two porosity proxy parameters among sites, S/V ratio (H = 11.03, p = 0.012) and percentage object volume (H = 10.41, p = 0.015). Dunn’s post-hoc tests with Bonferroni correction showed that Proximal Reef had a significantly lower S/V ratio than Mangrove Channel (p = 0.034) and Mangrove Lagoon (p = 0.027), and a significantly higher percentage object volume than Mangrove Channel (p = 0.016). No other pairwise comparisons among sites were significant (all adjusted p ≥ 0.086; Figure 3; Table S4). Beyond these two parameters, Kruskal-Wallis tests were used to test for overall differences among the four sites for each of the remaining morphometrics parameters at baseline (T1). These tests were also significant for five parameters: structure thickness (H = 11.33, p = 0.010), standard deviation of structure thickness (H = 10.83, p = 0.013), structure separation (H = 8.93, p = 0.030), intersection surface area (H = 8.78, p = 0.032), and standard deviation of structure separation (H = 8.48, p = 0.037; Tables S2 and S3). Baseline (T1) measurements showed that Proximal Reef samples had the densest skeletons with mean S/V ratio of 1.96 ± 0.67 mm⁻¹ and percentage object volume of 18.80 ± 4.71%, whereas Mangrove Lagoon and Mangrove Channel had the most porous skeletons with S/V ratio of 7.52 ± 2.29 and 7.00 ± 0.85 mm⁻¹ and percentage object volume 7.26 ± 3.04% and 5.88 ± 1.35%, respectively. Distal Reef samples had intermediate values for both S/V ratio (4.23 ± 1.13 mm⁻¹) and percentage object volume (10.96 ± 3.46%).

**Figure 3.**
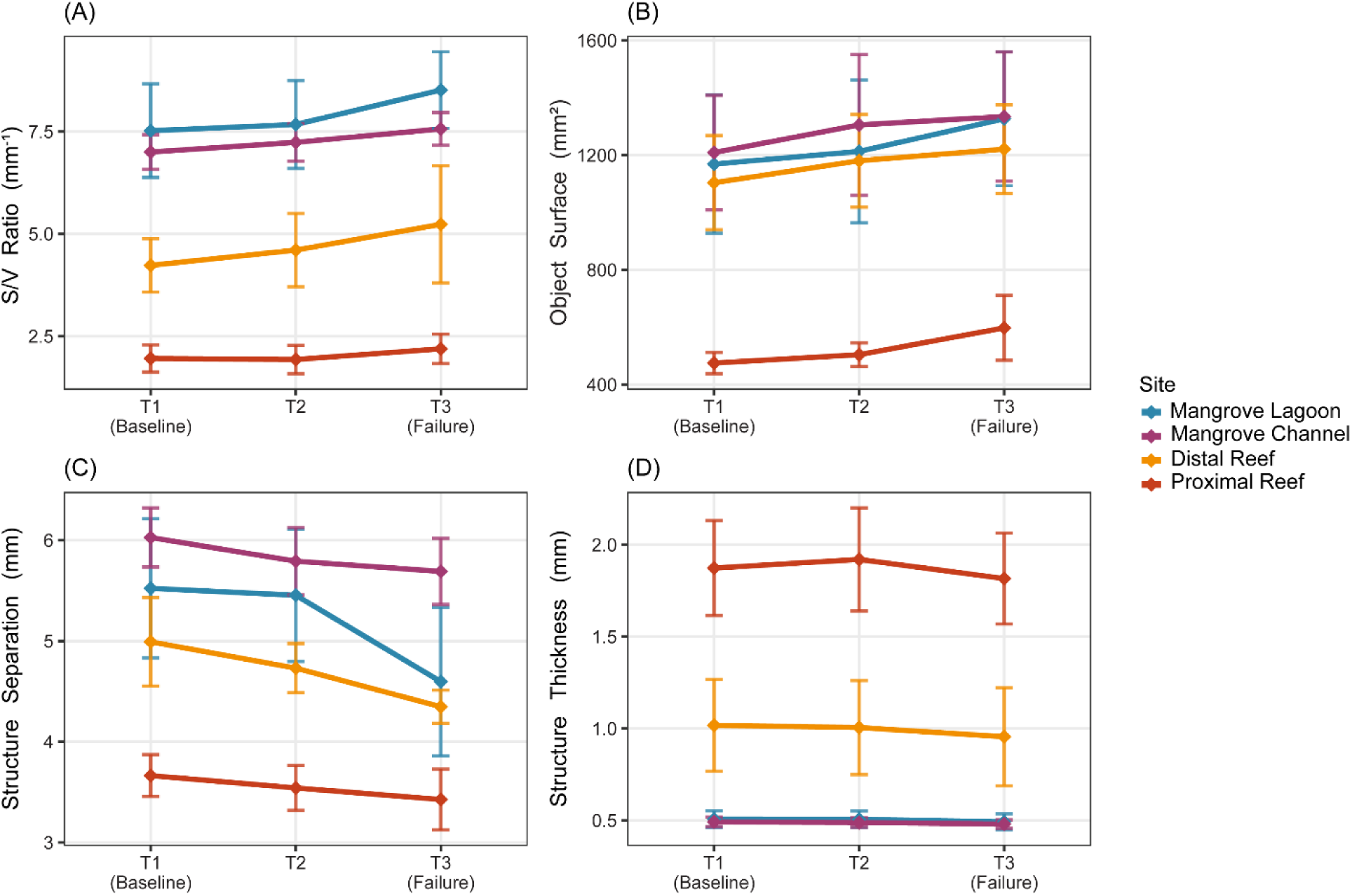
Longitudinal trajectories of four micro-CT morphometric parameters across the three compression time points (T1 = baseline, T2 = intermediate scan before failure, T3 = failure scan). (A) S/V ratio (mm⁻¹), (B) object surface area (mm²), (C) structure separation (mm), (D) structure thickness (mm). Lines represent site means ± SE. Colours indicate site. Note that T2 does not represent an equivalent loading state across samples (see Methods).

### Compression induces consistent microstructural changes

Progressive compression from T1 to T3 produced systematic and directionally consistent changes in micro-CT morphometric parameters across all 15 samples (Figure 4). Object surface area and surface density increased from T1 to T3 in all 15 samples with a mean of +16.1%, ranging from +2.6 to +63.8% (Wilcoxon p < 0.001). Structure separation decreased in all 15 samples with a mean of −10.4%, ranging between −33.4 to −1.1% (Wilcoxon p < 0.001). Intersection surface area declined in 13 of 15 samples (mean −33.1%; Wilcoxon p = 0.005). Structure linear density increased in 14 of 15 samples (mean +6.1%; Wilcoxon p < 0.001) and S/V ratio in 13 of 15 (mean +14.1%; Wilcoxon p < 0.001). Object volume and percentage object volume showed a modest increase in 13 of 15 samples with a mean of +2.1% (Wilcoxon p = 0.026 and 0.010, respectively).

**Figure 4.**
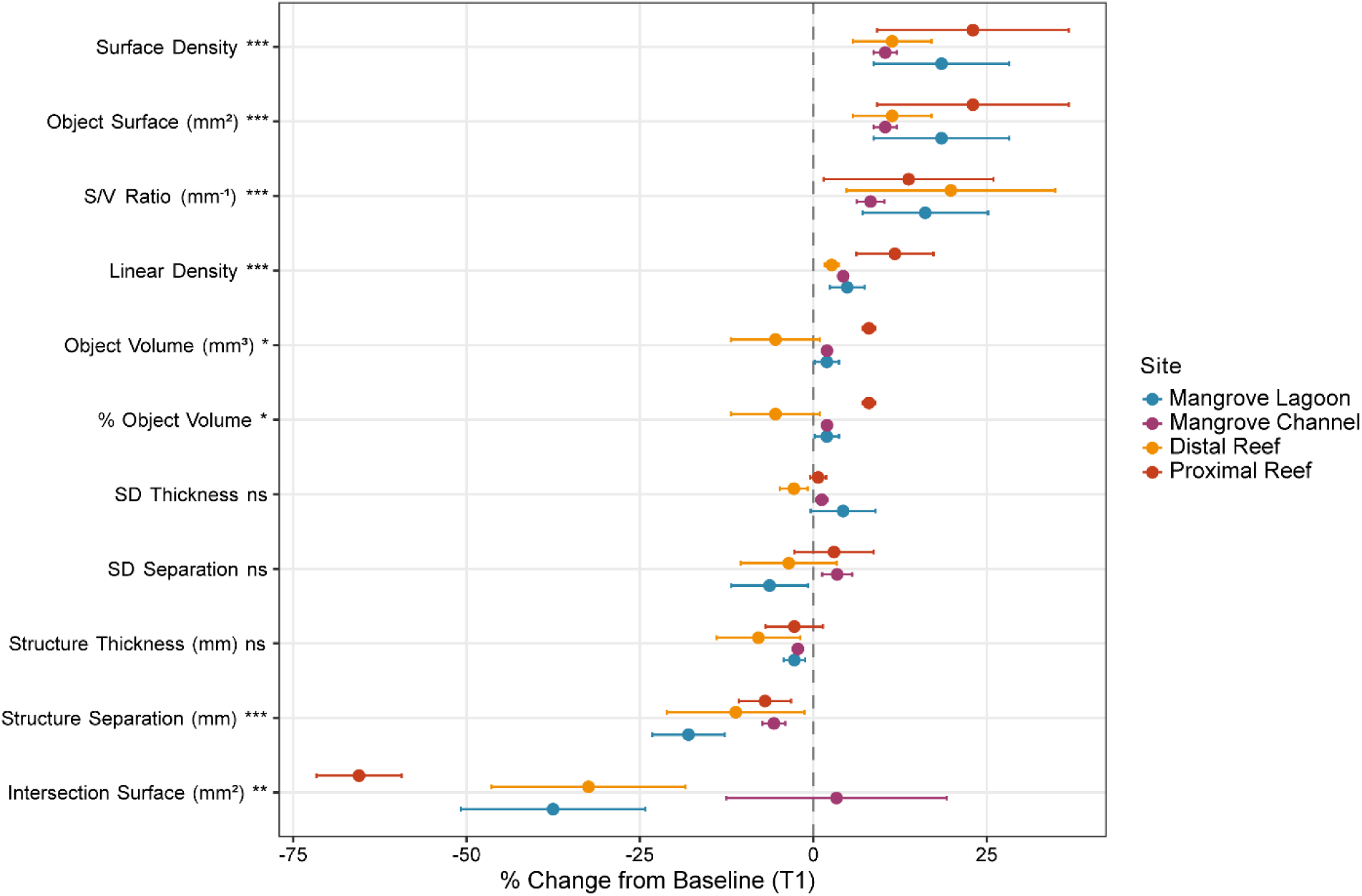
Mean percentage change from T1 to T3 for all eleven morphometric parameters. Points represent site means ± SE. Dashed reference line at zero denotes no change. Parameters are ordered by the pooled mean percentage change. Wilcoxon signed-rank significance: *** p < 0.001, ** p < 0.01, * p < 0.05, ns = not significant.

Structure thickness showed a consistent downward trend (−3.6%, 12 of 15 samples) that was not statistically significant (Wilcoxon p = 0.073). Standard deviations of both structure separation and structure thickness were not statistically different (Wilcoxon p = 0.978 and 0.720, respectively).

### Mechanical properties

Peak recorded forces ranged from 237.2 N (Mangrove Lagoon) to ≥3331.2 N (Proximal Reef; Figures 5 and 6). Proximal Reef samples consistently achieved the highest forces reaching 2015.95 N at confirmed brittle failure, with some samples exceeding 3331.2 N and 2113.0 N. All Mangrove Lagoon, Mangrove Channel, and Distal Reef samples remained below 730 N.

**Figure 5.**
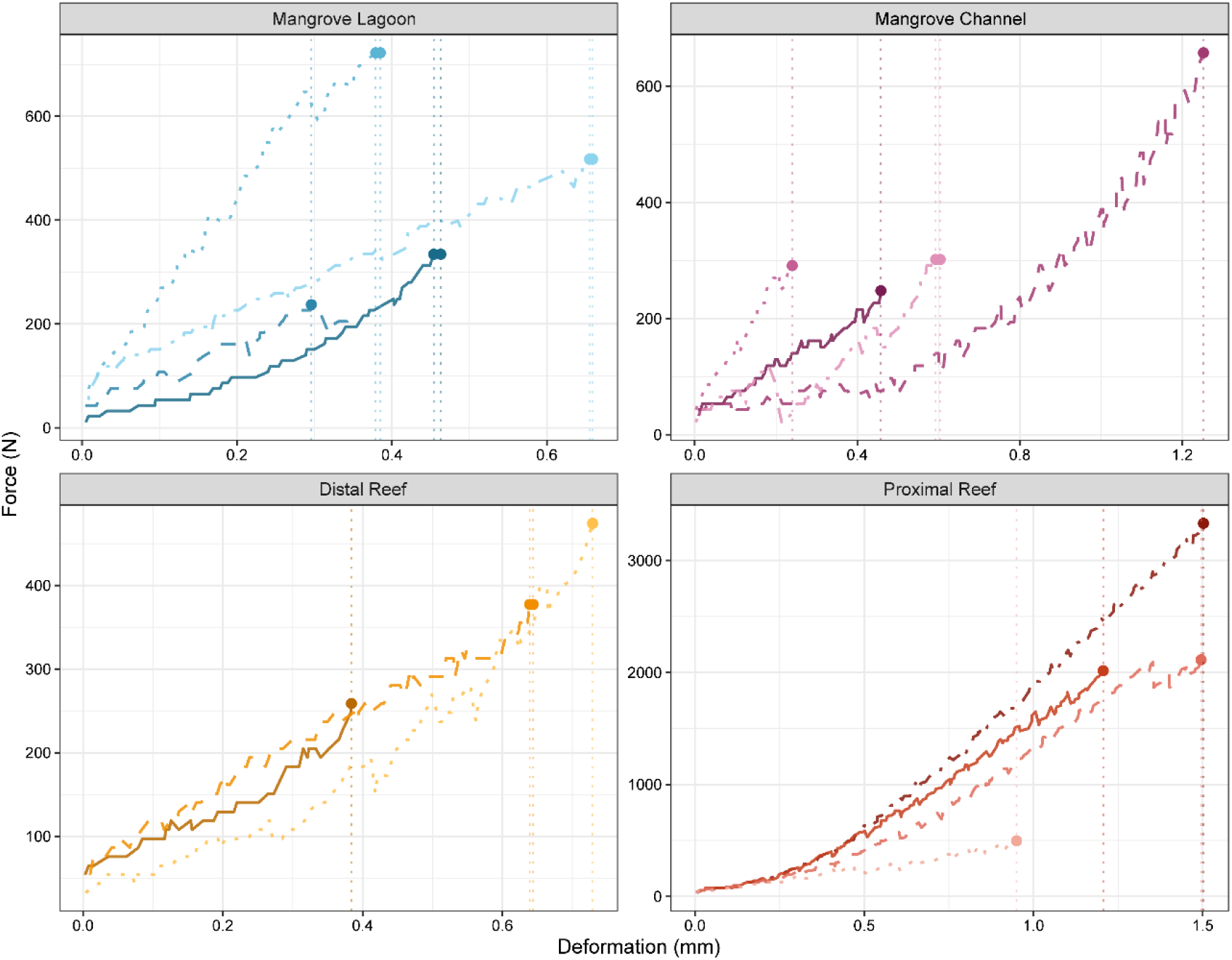
Force-displacement curves for all 15 samples grouped by site. Each curve shows force (N) against compressive deformation (mm) through the loading phase. Filled circles mark peak recorded force and dotted vertical lines indicate deformation at peak. Line style distinguishes individual samples.

**Figure 6.**
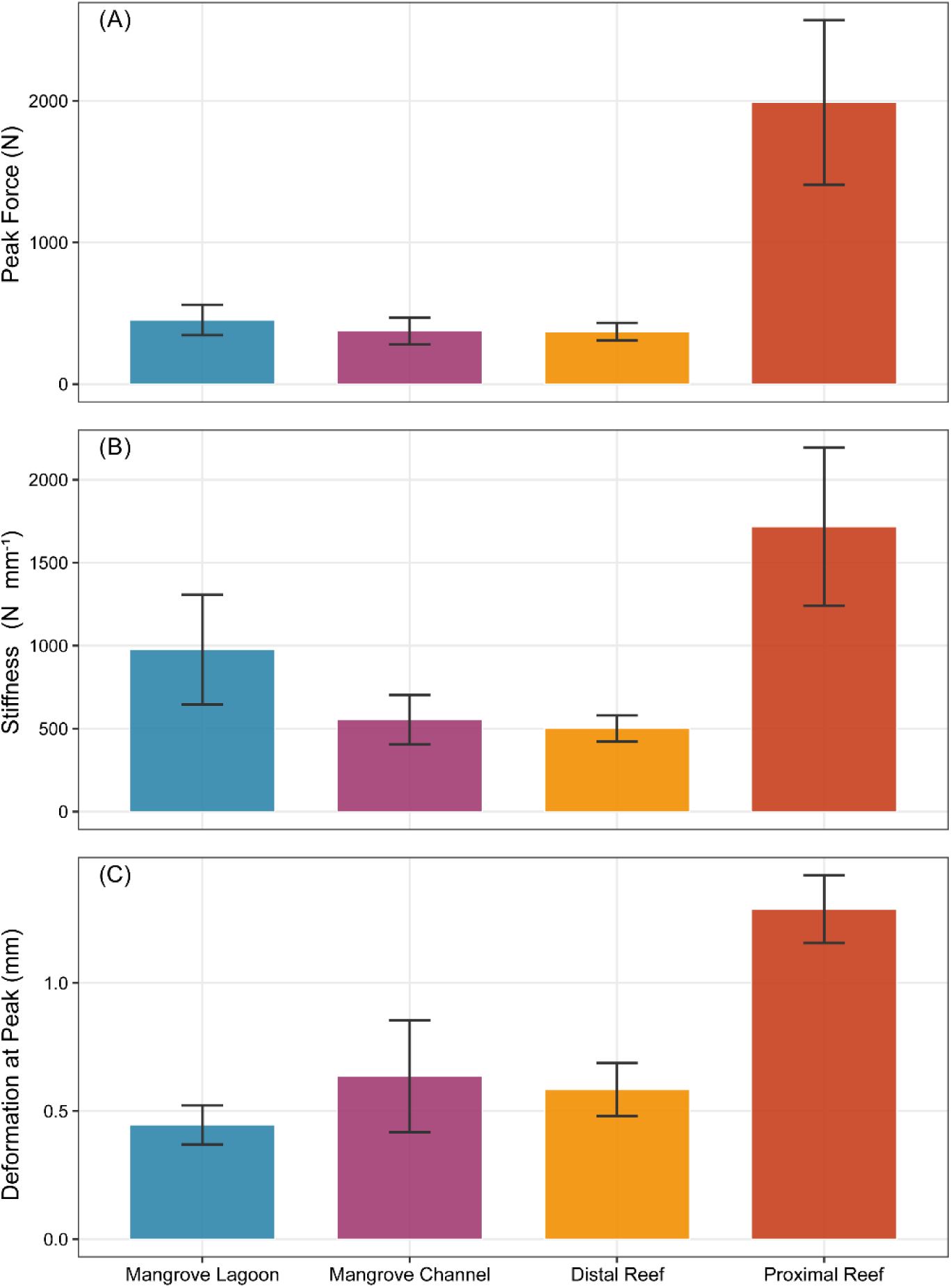
Mechanical properties by specimen and site. (A) Peak recorded force (N), (B) Stiffness (N mm⁻¹) and (C) Deformation at peak force (mm).

Stiffness ranged from 261.6 N mm⁻¹ (Mangrove Channel) to 2634.3 N mm⁻¹ (Proximal Reef). Site-level mean stiffness followed the same rank order as skeletal density. Proximal Reef was stiffest (1717.5 ± 951.1 N mm⁻¹), followed by Mangrove Lagoon (976.9 ± 659.8 N mm⁻¹), Mangrove Channel (554.2 ± 299.5 N mm⁻¹), and Distal Reef (501.3 ± 136.7 N mm⁻¹). This trend was not statistically significant (Kruskal-Wallis H = 3.65, p = 0.301), attributable to substantial within-site variance, particularly in Mangrove Lagoon (range 565-1962 N mm⁻¹) and Proximal Reef (range 382-2634 N mm⁻¹).

Ten of 15 samples had R² < 0.98 for the linear elastic region regression (Table S1). This reflects the fact that Mangrove Lagoon, Mangrove Channel, and Distal Reef samples tended to fail gradually, in a series of small steps rather than smoothly, with each step corresponding to a mini collapse of a skeletal element as loading was repeatedly paused and resumed for micro-CT scanning. All four Proximal Reef samples achieved R² ≥0.987, consistent with their denser and structurally more homogeneous skeletons producing cleaner elastic behaviour.

### Baseline porosity predicts stiffness

Baseline S/V ratio was negatively correlated with stiffness (Pearson r = −0.593, p = 0.020; Spearman ρ = −0.496, p = 0.060). Percentage object volume was positively correlated with stiffness (Pearson r = 0.648, p = 0.009; Spearman ρ = 0.457, p = 0.087). Across all eleven baseline (T1) morphometric parameters (Table 1), eight were significantly correlated with stiffness after Benjamini-Hochberg correction (all adjusted p < 0.05; Table S5), with the strongest correlations observed for the standard deviation of structure separation (Pearson r = −0.688, adjusted p = 0.019), object surface area (r = −0.681, adjusted p = 0.019), and structure thickness (r = 0.680, adjusted p = 0.019), followed by percentage object volume (r = 0.645, adjusted p = 0.022), intersection surface area (r = 0.642, adjusted p = 0.022), structure separation (r = −0.607, adjusted p = 0.028), the standard deviation of structure thickness (r = 0.594, adjusted p = 0.028), and S/V ratio (r = −0.592, adjusted p = 0.028). Object surface density, object volume, and structure linear density were not significantly correlated with stiffness (all adjusted p > 0.29). Pearson correlations were significant for the two focal porosity proxies while Spearman correlations were weaker and non-significant, indicating that the linear relationship was partly leveraged by Proximal Reef samples occupying extreme positions on both axes.

### Failure modes and microstructural deformation

Visual inspection of sequential micro-CT reconstructions revealed qualitatively distinct failure modes across sites, consistent with the differences in skeletal architecture and mechanical response described above (Figure 7).

**Figure 7.**
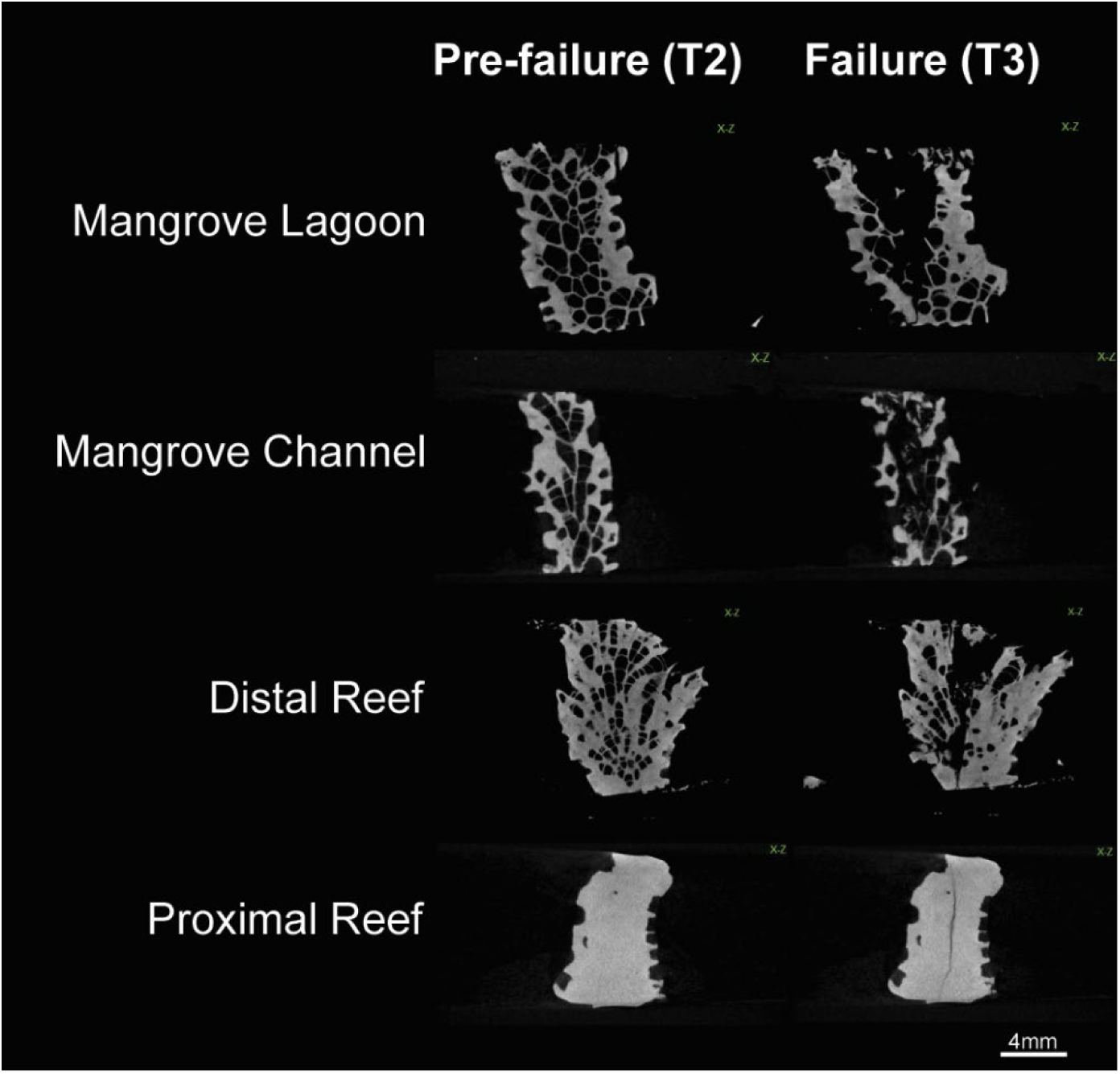
Representative failure modes of *Pocillopora* sp. skeletons under compression across the four sites. Paired pre-failure (T2) and post-failure (T3) micro-CT reconstructions (X–Z) are shown for one representative sample from each site. Force (N) and displacement (mm) at T3 are indicated on the right.

In Mangrove Lagoon and Mangrove Channel samples, pre-failure scans (T2) showed highly open skeletal strut architectures with large inter-skeletal pore spaces and slender branching elements. Post-failure (T3), these samples exhibited extensive fragmentation, with the porous skeletal framework collapsing and separating into disconnected debris. This failure was accompanied by lateral redistribution of skeletal material, and transverse (X–Y) sections revealed disintegration of the original circular specimen cross-section into multiple displaced fragments. The degree of structural disintegration was pronounced by large portions of the specimen that were displaced well outside the original footprint (Figure 7; Mangrove Lagoon).

Distal Reef samples show an intermediate failure mode. Baseline architectures were denser than Mangrove Lagoon and Mangrove Channel but retained distinct porous organisation. Post-failure scans showed partial compaction where the skeletal mass was compressed and somewhat consolidated rather than explosively fragmented, though crack planes were visible in the longitudinal (X-Z) view and some lateral displacement of large fragments occurred (Figure 6; Distal Reef).

Proximal Reef samples exhibited a qualitatively different failure mode reflecting their dense, low-porosity architecture. Pre-failure, these samples presented as near-solid blocks with very few visible pores and thick, well-connected skeletal elements (Figure 7). Post-failure, the predominant damage pattern is the propagation of discrete crack planes through the specimen rather than generalised fragmentation. For example, in one sample, a vertical crack plane was clearly visible in the X-Z section, while the overall specimen geometry remained largely intact. In another sample, the X-Y section revealed a primary fracture splitting the circular cross-section into two major lobes, with minimal debris.

## Discussion

### Skeletal density and compressive performance reflect habitat differences

Our results demonstrate a consistent change in both skeletal architecture and mechanical performance across the mangrove to reef habitat gradient at Low Isles. Proximal Reef samples possessed the least porous skeletons with the highest compressive stiffness and peak forces with values two to three times greater than those recorded at the two mangrove sites and Distal Reef. While wave exposure differs across sites, this pattern is unlikely to be driven by wave exposure alone and may be more strongly influenced by water quality at the mangrove system (Chadda-Harmer et al., 2025b). Wave direction at Low Isles is predominantly from the south-east, leaving Proximal Reef relatively sheltered from wave action by Woody Island and an adjacent sandbar, whereas Distal Reef lies along the more wave-exposed outer reef margin (Schueth and Frank, 2008). Proximal Reef serves as the key tidal channel for Low Isles, suggesting that tidal flow may be the dominant hydrodynamic influence at this site (Schueth and Frank, 2008). Previous work from the same sites investigating *Acropora millepora* colonies also found that Proximal Reef had less porous skeletons than those at Distal Reef (Chadda-Harmer et al., 2025a). Notably, despite being the most wave exposed of the four sites, Distal Reef did not have the highest bulk density or stiffness, further indicating that wave exposure alone does not explain the observed gradient.

Similarly, FORAM Index-based water quality assessments across the Low Isles system showed that suitability for coral growth did not scale simply with proximity to mangroves or wave exposure but instead depended on hydrodynamic forcing and geomorphological setting, including the position of tidal conduits (Chadda-Harmer et al., 2025b). As the main conduit, Proximal Reef may receive more water exchange from outside the mangrove system, potentially reducing exposure to mangrove-derived stress relative to Distal Reef. Together, these observations suggest that tidal flow mediated water quality, in addition to wave-driven mechanical selection, may contribute to the skeletal density and stiffness gradient observed here. This is consistent with the general relationship between hydrodynamic exposure and skeletal density in corals, whereby higher water flow can enhance calcification through improved nutrient delivery while also imposing direct mechanical selection on structural robustness (Comeau et al., 2014; Madin and Connolly, 2006).

Microstructural porosity may also be relatively insensitive to moderate differences in wave exposure. For example, a micro-CT study of *Acropora pulchra* across a wave-exposed-to-sheltered reef-flat gradient found no significant difference in skeletal porosity despite the exposure gradient (Roche et al., 2010). This suggests that microarchitecture may be less affected by wave exposure than external morphology, further supporting that the differences observed among our sites are due to differences in water quality. The increase in skeletal density from the Mangrove Lagoon to the Proximal Reef broadly matches zonation patterns reported across wave-exposure gradients in other reef systems (Doszpot et al., 2019; Madin et al., 2014) and is consistent with Proximal Reef’s status as the least mangrove influenced site of the four in terms of tidal water exchange (Chadda-Harmer et al., 2025a). Importantly, our findings directly quantify compressive mechanical performance, demonstrating that habitat-related differences in skeletal architecture translate into substantial differences in structural resistance.

There was a stronger relationship between stiffness and percentage object volume, than stiffness and S/V ratio, which is mechanically informative. This strong correlation between percentage object volume and stiffness suggests that the overall amount of skeletal material is a meaningful predictor of compressive resistance for these samples. While S/V quantifies the extent and complexity of surface-connected architecture, percentage object volume more directly reflects the fraction of solid material available to bear compressive loads (Gibson and Ashby, 2010; Keaveny et al., 2001). Mechanical performance in porous biological materials is governed not only by overall density but also by the thickness, connectivity, and spatial arrangement of load bearing elements (Keaveny et al., 2001; Wang et al., 2015). Consequently, coral skeletons with a relatively high S/V ratio may still maintain substantial mechanical resistance if the load-bearing skeletal struts remain thick and well connected. Studies of skeletal systems demonstrate that elastic modulus and strength align strongly with skeletal strut morphology and connectivity (Keaveny et al., 2001; Wang et al., 2015). Our results suggest that bulk skeletal volume fraction is therefore a more direct predictor of compressive performance than S/V alone, while also demonstrating the value of combining volumetric and architectural measures when assessing coral skeletal mechanics. Environmental conditions can influence both skeletal structure and mechanical behaviour in corals (Mollica et al., 2018; Moynihan et al., 2021). This finding highlights a methodological limitation of relying solely on structural metrics as a proxy for skeletal hardness or strength and demonstrates the value of incorporating volumetric measures of skeletal architecture when assessing coral mechanical performance (Chadda-Harmer et al., 2025a; Chamberlain, 1978; Mwachireya et al., 2016).

Correlations between stiffness and the eleven baseline morphometric parameters reinforce this mechanistic interpretation. Parameters describing strut dimension and spatial arrangement such as structure thickness, the variability in structure separation and object surface area, were consistently the strongest correlates of stiffness, consistent with cellular solids theory, in which strut geometry is a primary determinant of bulk mechanical resistance (Gibson and Ashby, 2010). In contrast, parameters describing sheer material quantity such as object volume, structure linear density, and object surface density showed weak, non-significant relationships with stiffness, indicating that raw material quantity or topological branching characteristics alone are insufficient predictors of compressive resistance. Together, these results suggest that skeletal parameters reflecting load-bearing strut geometry and spatial organisation, rather than bulk material presence per se, are more informative predictors of compressive stiffness.

Site-level stiffness followed the density gradient across all four sites, from lowest at Distal Reef and Mangrove Channel, through Mangrove Lagoon, to the highest at Proximal Reef. Despite the directional trend, the between-site stiffness difference was not statistically significant, in contrast to the significant between-site differences in skeletal architecture identified by Chadda-Harmer et al. 2025a. This likely reflects limited statistical power, as sample sizes were constrained to n = 3-4 per site and within-site variance was substantial, particularly in Mangrove Lagoon and Proximal Reef. Under these conditions, only very large effect sizes are detectable. The observed within-site variability likely represents genuine biological variation in skeletal architecture at the colony level, which is well documented in *Pocillopora* and other branching corals, where morphology and skeletal traits can vary strongly in response to local environmental conditions and phenotypic plasticity (Doszpot et al., 2019; Paz-García et al., 2015; Todd, 2008).

Across the sites, skeletal stiffness generally increased with increasing skeletal density and decreasing porosity, with the strongest values occurring at Proximal Reef, which also had the highest relative skeletal density and lowest porosity. Pearson correlations indicated significant positive associations between stiffness and measures of skeletal density and significant negative associations with porosity. In contrast, the corresponding Spearman rank correlations were weaker and non-significant. Because rank-order analysis is comparatively less influenced by extreme values, the weaker Spearman correlation suggests that the Proximal Reef samples, which occupy the upper extremes of both skeletal density and stiffness, contribute substantially to this relationship. This pattern suggests that the relationship between skeletal porosity and mechanical performance may be non-linear, likely reflecting the power-law scaling relationships widely described for cellular and porous materials, in which stiffness and strength scale with relative density (Gibson and Ashby, 2010; Keaveny et al., 2001). The apparent linear trend observed here may partially be due to the separation between the Proximal Reef cluster and the remaining sites, rather than representing a universal linear relationship across the full range of skeletal architectures.

### Failure mode reflects skeletal architecture

Qualitative assessments of failure mode across the coral fragments revealed that skeletal architecture influenced not only the magnitude of mechanical resistance but also the pathway of structural failure. In Mangrove Lagoon and Mangrove Channel samples, the open, porous architecture provided numerous crack initiation sites but limited structural redundancy. Consequently, once critical skeletal struts failed, alternative load paths are limited, and cascading collapse of the thin skeletal network resulted in rapid fragmentation and loss of structural coherence. In contrast, Proximal Reef samples exhibited failure through the formation of discrete crack planes within an otherwise intact skeleton. This crack-plane failure mode is characteristic of dense porous materials, in which a high mineral volume fraction and greater connectivity among load bearing elements permit stress redistribution around propagating cracks, delaying catastrophic collapse and maintaining structural integrity over a greater deformation range (Gibson and Ashby, 2010). Rather than fragmenting into disconnected debris, the dense skeletal framework remains coherent until failure localises along a distinct fracture plane. This contrast highlights how differences in skeletal architecture can fundamentally alter not only the strength but also the failure trajectory of coral skeletons under compressive loading, consistent with established behaviour in cellular and porous materials (Gibson and Ashby, 2010; Wang et al., 2015). This distinction has direct relevance to breakage under wave action or cyclone impacts in the field. A dense, structurally continuous skeleton is more likely to withstand a single high-energy loading event intact, whereas a porous skeleton that fails by fragmentation is more susceptible to shattering into multiple pieces under storm-driven wave forces (Madin et al., 2014; Madin and Connolly, 2006).

These contrasting failure modes have implications beyond ultimate failure force. Fragmentation-dominated failure generates loose skeletal debris and results in a near-complete loss of the colony’s structural contribution to the reef framework, reducing habitat complexity and weakening the physical integrity of the reef matrix (Alvarez-Filip et al., 2009; Graham and Nash, 2012; Richardson et al., 2020). In contrast, crack-plane failure may preserve partial load bearing capacity and retain substrate suitable for recovery or recolonisation (Madin et al., 2014; Perry et al., 2018). Both failure modes ultimately result in colony damage, however, their consequences for reef framework persistence and recovery trajectories may differ. Fragmentation causes reef framework breakdown and rubble formation, whereas more coherent failure can preserve some structural function, affecting recovery after disturbance and long-term reef resilience (Madin and Connolly, 2006; Perry et al., 2018).

The force-displacement curves are consistent with these contrasting failure modes. Samples from the two mangrove sites exhibited irregular, stepped loading profiles indicative of sequential micro-failure events, in which individual skeletal elements collapsed progressively under increasing load. Distal Reef samples showed an intermediate pattern, with fewer and less pronounced stepped events than the two mangrove sites, consistent with their intermediate skeletal density and partial structural collapse failure mode. In contrast, Proximal Reef samples displayed smoother, more continuous loading curves prior to peak force, reflecting a more homogeneous distribution of stress through a denser and better connected skeletal network. These differences in failure behaviour are also reflected in the lower R² values obtained for the linear elastic region in the Mangrove Lagoon, Mangrove Channel, and Distal Reef samples, where progressive sub-failure damage obscured the definition of a distinct elastic phase.

Importantly, despite substantial differences in skeletal density, stiffness, and ultimate failure force among sites, the direction of microstructural change during compression was broadly consistent across all sites. The increase in object surface area and surface density indicates the formation of new mineral surfaces during compression-induced microcracking, consistent with micro-CT observations of compressed coral reef limestone (Meng et al., 2022). At the same time, the decrease in structure separation indicates progressive compaction of pore space as skeletal elements were displaced under load. The decline in intersection surface area is consistent with disconnection and loss of connectivity among skeletal elements as fracture propagated through the network, and the increases in structure linear density and S/V ratio are both consistent with a rise in fractured surface area per unit volume. Together, these changes describe a coherent degradation pathway characterised by surface generating fracture, strut compaction, and progressive network disconnection, broadly comparable to the pore-structure-dependent fracture pathways described for coral reef limestone under compressive loading (Xu et al., 2025). The non-significant decline in structure thickness further suggests that crushing and shortening of individual skeletal struts occurred during compression, although the extent of this damage varied among specimens. Interestingly, despite pronounced shifts in mean architectural properties, the within-specimen variability of structure thickness and separation remained largely unchanged, indicating that compression altered the average structure of the skeletal network without systematically changing its spatial heterogeneity. These findings suggest that while habitat influences the magnitude of mechanical resistance and the mode of ultimate failure, the underlying processes governing compression induced skeletal damage are broadly conserved across habitats.

### Implications for mangrove associated coral reefs

The lower stiffness and fragmentation prone failure mode of mangrove site samples suggests that physiological tolerance alone may not predict resilience to future environmental change. Mangrove associated corals persist under stressful environmental conditions, including elevated temperatures, variable and often low pH, and fluctuating oxygen, that cause bleaching and mortality elsewhere (Camp et al., 2019, 2018; Yates et al., 2014). Their persistence is therefore often interpreted as evidence of pre-adaptation to future ocean conditions (Burt et al., 2020; Scucchia et al., 2023). However, our results suggest that this physiological tolerance come at the cost of reduced skeletal mechanical performance relative to open-reef counterparts.

This trade-off may reflect fundamentally different selective environments. In the low-energy, hydrodynamically protected Mangrove Lagoon, investment in maximum skeletal hardness confers little advantage since physical disturbance from wave action is rare, and resources may be more advantageously allocated to stress tolerance, rapid growth, or reproductive output (Camp et al., 2018; Palumbi et al., 2014; Scucchia et al., 2023). Conversely, in the Proximal Reef environment, structural reinforcement is critical for survival and is strongly selected. Further, despite the Mangrove Channel also possessing a strong tidal flow (Chadda-Harmer et al., 2025a), corals here possessed similar structural properties to the lagoon site, highlighting the potential influence of water quality conditions amongst the mangroves beyond a purely hydrodynamic lens. Likewise, Distal Reef, despite its position on the more wave-exposed outer reef margin, showed only intermediate skeletal density and stiffness, consistent with its intermediate mangrove influence and suggesting that resource allocation to skeletal reinforcement at this site is shaped by more than just wave exposure alone. This resource allocation framework, in which different environmental pressures favour different aspects of skeletal investment, is consistent with the adaptive plasticity hypothesis proposed for corals across environmental gradients (Palumbi et al., 2014; Todd, 2008).

Importantly, if climate change increases the intensity or frequency of physical disturbances in currently protected habitats, through the penetration of storm surge into nearshore mangrove environments, or through shifts in the trajectories of intensifying cyclones, mangrove associated corals may face novel selective pressures for which their skeletal mechanical properties provide limited resilience (Baldock et al., 2014; Burt et al., 2020; IPCC, 2023; Scucchia et al., 2023). These results have a broader implication for corals persisting under suboptimal or highly variable conditions. The same environmental variability that may favour physiological stress tolerance could simultaneously compromise the skeletal mechanical properties required to withstand acute physical disturbance. Thus, tolerance to one stressor should not be assumed to confer resilience to other, potentially interacting stressors. Because mangrove lagoons are increasingly used as natural analogues of future ocean conditions (Camp et al., 2019; Scucchia et al., 2023), this trade-off suggests that corals acclimatised to warmer, more acidic, and hypoxic waters may face a future characterised by both chronic environmental stress and intensifying storm activity, while possessing skeletons that are structurally less capable of resisting fragmentation under physical loading. While environments characterised by fluctuating water chemistry may enhance coral resilience to future ocean conditions and provide potential sources of climate-resilient corals for restoration (e.g. Morikawa and Palumbi 2019), it will be important to determine whether this resilience comes at the cost of compromised skeletal mechanical performance.

### Limitations and future directions

Several limitations constrain the interpretation of the present results. Sample sizes of n = 3-4 per site provide limited statistical power, and the destructive nature of compression testing prevents replication at the colony level. The in-situ micro-CT compression protocol requires small sample sections (approximately 1 cm), which may not represent the full range of skeletal architecture across a colony and cannot capture whole colony structural properties such as branch thickness or branching geometry. Future work should combine micro-scale compression data with macro-scale structural assessments.

Integration of environmental data, particularly hydrodynamic exposure histories, thermal anomaly records, and carbonate chemistry for each site, is needed to directly link the skeletal property gradients observed here to their environmental drivers. Calcification rate data and stable isotope analyses (δ¹³C, Sr/Ca) would further illuminate the relationships among growth rate, skeletal architecture, crystal organisation, and the mechanical properties reported here. Importantly, characterising hydrodynamic conditions alongside water-quality parameters would help disentangle the relative drivers of variation in skeletal properties.

## Conclusions

This study demonstrates that *Pocillopora* sp. corals across a mangrove to reef gradient at Low Isles, GBR, exhibit substantial differences in skeletal architecture, mechanical performance, and failure mode that reflect the mechanical selective environment of each habitat. Proximal Reef samples were two to three times stiffer and withstood two to three times greater peak forces than other sites samples and fail through discrete crack propagation rather than the skeletal network fragmentation observed in Mangrove Lagoon and Mangrove Channel samples. Distal Reef samples occupied an intermediate position in both skeletal density and stiffness despite their outer-reef, wave-exposed setting, underscoring that tidal-flow-mediated water quality, rather than wave exposure alone, shapes this gradient. Bulk mineral density (percentage object volume) is a stronger predictor of stiffness than skeletal architectural complexity (S/V ratio), highlighting the importance of distinguishing between different structural metrics when interpreting the mechanical consequences of skeletal architecture. Compression consistently increased surface area, decreased structure separation, and reduced skeletal connectivity across all sites, irrespective of initial skeletal density, describing a universal pattern of mechanical degradation. These findings underscore that assessing the future resilience of mangrove associated coral populations, as well as corals from other marginal and extreme environments, must extend beyond physiological stress tolerance to include the structural mechanical properties that govern resistance to physical pressures under intensifying climate driven environmental change.

## Author contribution

DCH and SF conceived the overarching study; SF designed the field sampling framework; AZ and MF developed the compression experimental methodology, conducted the experiments and collected the data; AZ analysed the data and led the writing of the manuscript; DCH, SF and MF edited and reviewed the manuscript. All authors contributed critically to the drafts and gave final approval for publication.

## Data availability statement

All relevant data and details of resources can be found within the article and its supplementary information.

## References

Al-Horani, F.A., 2015. Physiology of Skeletogenesis in Scleractinian Coral, in: Diseases of Coral. John Wiley and Sons Inc, pp. 192–205. 10.1002/9781118828502.ch12

Altintas, G., Cankal, D., Kilic, G., Ergun, S., 2024. Mechanical properties of Acropora cervicornis aragonite skeleton by using multiscale models based on micro-CT data. Advances in Applied Ceramics 123, 59–66. 10.1177/17436753241276985

Alvarez-Filip, L., Dulvy, N.K., Gill, J.A., Côté, I.M., Watkinson, A.R., 2009. Flattening of Caribbean coral reefs: Region-wide declines in architectural complexity. Proceedings of the Royal Society B: Biological Sciences 276, 3019–3025. 10.1098/rspb.2009.0339

Anthony, K.R.N., 2016. Coral Reefs under Climate Change and Ocean Acidification: Challenges and Opportunities for Management and Policy. Annu. Rev. Environ. Resour. 41, 59–81. 10.1146/annurev-environ-110615-085610

Baldock, T.E., Karampour, H., Sleep, R., Vyltla, A., Albermani, F., Golshani, A., Callaghan, D.P., Roff, G., Mumby, P.J., 2014. Resilience of branching and massive corals to wave loading under sea level rise - A coupled computational fluid dynamics-structural analysis. Mar. Pollut. Bull. 86, 91–101. 10.1016/j.marpolbul.2014.07.038

Bertini, L., Hendy, E., Summerfield, R., Fernandez, V., Clark, B., Martin-Silverstone, E., Lucka, F., Kiss, M.B., Johnson, K., 2026. How to quantify and minimise error in coral skeletal density estimates using X-ray µCT. R. Soc. Open Sci. 13. 10.1098/RSOS.251288

Burt, J.A., Camp, E.F., Enochs, I.C., Johansen, J.L., Morgan, K.M., Riegl, B., Hoey, A.S., 2020. Insights from extreme coral reefs in a changing world. Coral Reefs. 10.1007/s00338-020-01966-y

Camp, E.F., Edmondson, J., Doheny, A., Rumney, J., Grima, A.J., Huete, A., Suggett, D.J., 2019. Mangrove lagoons of the Great Barrier Reef support coral populations persisting under extreme environmental conditions. Mar. Ecol. Prog. Ser. 625, 1–14. 10.3354/MEPS13073

Camp, E.F., Nitschke, M.R., Rodolfo-Metalpa, R., Houlbreque, F., Gardner, S.G., Smith, D.J., Zampighi, M., Suggett, D.J., 2017. Reef-building corals thrive within hot-acidified and deoxygenated waters. Sci. Rep. 7, 1–9. 10.1038/s41598-017-02383-y

Camp, E.F., Schoepf, V., Mumby, P.J., Hardtke, L.A., Rodolfo-Metalpa, R., Smith, D.J., Suggett, D.J., 2018. The future of coral reefs subject to rapid climate change: Lessons from natural extreme environments. Front. Mar. Sci. 10.3389/fmars.2018.00004

Capasso, L., Aranda, M., Cui, G., Pousse, M., Tambutté, S., Zoccola, D., 2022. Investigating calcification-related candidates in a non-symbiotic scleractinian coral, Tubastraea spp. Sci. Rep. 12. 10.1038/s41598-022-17022-4

Chadda-Harmer, D., Byrne, M., Foley, M., Barrington-Smith, G., Camp, E.F., Foo, S.A., 2025a. Skeletal structure and adaptive mechanisms of corals inhabiting a mangrove environment. Coral Reefs 1–18. 10.1007/S00338-025-02727-5/FIGURES/1

Chadda-Harmer, D., Byrne, M., Reymond, C.E., Fellowes, T.E., Camp, E.F., Foo, S.A., 2025b. Benthic foraminifera as bioindicators of coral condition near mangrove environments. Mar. Environ. Res. 209. 10.1016/j.marenvres.2025.107159

Chamberlain, J.A., 1978. Mechanical properties of coral skeleton: compressive strength and its adaptive significance. Paleobiology 4, 419–435. 10.1017/S0094837300006163

Cohen, A.L., Holcomb, M., 2009. Why Corals Care About Ocean Acidification: Uncovering the Mechanism 22, 118–127. 10.2307/24861029

Comeau, S., Edmunds, P.J., Lantz, C.A., Carpenter, R.C., 2014. Water flow modulates the response of coral reef communities to ocean acidification. Scientific Reports 2014 4:1 4, 6681-. 10.1038/srep06681

Coronado, I., Fine, M., Bosellini, F.R., Stolarski, J., 2019. Impact of ocean acidification on crystallographic vital effect of the coral skeleton. Nature Communications 10. 10.1038/s41467-019-10833-6

Davies, P.S., 1989. Short-term growth measurements of corals using an accurate buoyant weighing technique. Mar. Biol. 101, 389–395. 10.1007/BF00428135

Doszpot, N.E., McWilliam, M.J., Pratchett, M.S., Hoey, A.S., Figueira, W.F., 2019. Plasticity in three-dimensional geometry of branching corals along a cross-shelf gradient. Diversity (Basel). 11. 10.3390/D11030044

Falter, J.L., Lowe, R.J., Zhang, Z., McCulloch, M., 2013. Physical and Biological Controls on the Carbonate Chemistry of Coral Reef Waters: Effects of Metabolism, Wave Forcing, Sea Level, and Geomorphology. PLoS One 8, e53303. 10.1371/JOURNAL.PONE.0053303

Fantazzini, P., Mengoli, S., Pasquini, L., Bortolotti, V., Brizi, L., Mariani, M., Di Giosia, M., Fermani, S., Capaccioni, B., Caroselli, E., Prada, F., Zaccanti, F., Levy, O., Dubinsky, Z., Kaandorp, J.A., Konglerd, P., Hammel, J.U., Dauphin, Y., Cuif, J.P., Weaver, J.C., Fabricius, K.E., Wagermaier, W., Fratzl, P., Falini, G., Goffredo, S., 2015. Gains and losses of coral skeletal porosity changes with ocean acidification acclimation. Nature Communications 2015 6:1 6, 7785-. 10.1038/ncomms8785

Frank, T.D., Jell, J.S., 2006. Recent developments on a nearshore, terrigenous-influenced reef: Low Isles Reef, Australia. J. Coast. Res. 22, 474–486. 10.2112/03-0127.1

Gibson, L.J., 2003. Cellular Solids. MRS Bull. 28, 270–274. 10.1557/MRS2003.79

Gibson, L.J., Ashby, M.F., 2010. Cellular materials in nature and medicine. Integr. Comp. Biol. 51, E47--E47.

Graham, N.A.J., Nash, K.L., 2012. The importance of structural complexity in coral reef ecosystems. Coral Reefs 2012 32:2 32, 315–326. 10.1007/S00338-012-0984-Y

Hamylton, S.M., McLean, R., Lowe, M., Adnan, F.A.F., 2019. Ninety years of change on a low wooded island, Great Barrier Reef. R. Soc. Open Sci. 6. 10.1098/rsos.181314

Haydon, T.D., Matthews, J.L., Seymour, J.R., Raina, J.B., Seymour, J.E., Chartrand, K., Camp, E.F., Suggett, D.J., 2023. Metabolomic signatures of corals thriving across extreme reef habitats reveal strategies of heat stress tolerance. Proceedings of the Royal Society B: Biological Sciences 290. 10.1098/RSPB.2022.1877/86633

Haydon, T.D., Seymour, J.R., Raina, J.-B., Edmondson, J., Siboni, N., Matthews, J.L., Camp, E.F., Suggett, D.J., 2021. Rapid Shifts in Bacterial Communities and Homogeneity of Symbiodiniaceae in Colonies of Pocillopora acuta Transplanted Between Reef and Mangrove Environments. Front. Microbiol. 12, 3131. 10.3389/fmicb.2021.756091

Hoegh-Guldberg, O., Mumby, P.J., Hooten, A.J., Steneck, R.S., Greenfield, P., Gomez, E., Harvell, C.D., Sale, P.F., Edwards, A.J., Caldeira, K., Knowlton, N., Eakin, C.M., Iglesias-Prieto, R., Muthiga, N., Bradbury, R.H., Dubi, A., Hatziolos, M.E., 2007. Coral reefs under rapid climate change and ocean acidification. Science. 10.1126/science.1152509

IPCC, 2023. Climate Change 2023: Synthesis Report. Contribution of Working Groups I, II and III to the Sixth Assessment Report of the Intergovernmental Panel on Climate Change [Core Writing Team, H. Lee and J. Romero (eds.)]. IPCC, Geneva, Switzerland. 10.59327/IPCC/AR6-9789291691647

Keaveny, T.M., Morgan, E.F., Niebur, G.L., Yeh, O.C., 2001. Biomechanics of trabecular bone. Annu. Rev. Biomed. Eng. 3, 307–333. 10.1146/ANNUREV.BIOENG.3.1.307/CITE/REFWORKS

Madin, J.S., Baird, A.H., Dornelas, M., Connolly, S.R., 2014. Mechanical vulnerability explains size-dependent mortality of reef corals. Ecol. Lett. 17, 1008–1015. 10.1111/ele.12306

Madin, J.S., Connolly, S.R., 2006. Ecological consequences of major hydrodynamic disturbances on coral reefs. Nature 2006 444:7118 444, 477–480. 10.1038/nature05328

Madin, J.S., O’Donnell, M.J., Connolly, S.R., 2008. Climate-mediated mechanical changes to post-disturbance coral assemblages. Biol. Lett. 4, 490–493. 10.1098/rsbl.2008.0249

Meng, Q., Wu, K., Zhou, H., Qin, Q., Wang, C., 2022. Mesoscopic damage evolution of coral reef limestone based on real-time CT scanning. Eng. Geol. 307. 10.1016/j.enggeo.2022.106781

Milita, S., Zaquin, T., Fermani, S., Montroni, D., Pinkas, I., Barba, L., Falini, G., Mass, T., 2023. Assembly of the Intraskeletal Coral Organic Matrix during Calcium Carbonate Formation. Cryst. Growth Des. 23, 5801–5811. 10.1021/acs.cgd.3c00401

Mollica, N.R., Guo, W., Cohen, A.L., Huang, K.F., Foster, G.L., Donald, H.K., Solow, A.R., 2018. Ocean acidification affects coral growth by reducing skeletal density. Proc. Natl. Acad. Sci. U. S. A. 115, 1754–1759. 10.1073/pnas.1712806115

Morikawa, M.K., Palumbi, S.R., 2019. Using naturally occurring climate resilient corals to construct bleaching-resistant nurseries. Proceedings of the National Academy of Sciences 116, 10586–10591. 10.1073/pnas.1721415116

Moynihan, M.A., Amini, S., Goodkin, N.F., Tanzil, J.T.I., Chua, J.Q.I., Fabbro, G.N., Fan, T.Y., Schmidt, D.N., Miserez, A., 2021. Environmental impact on the mechanical properties of Porites spp. corals. Coral Reefs 40, 701–717. 10.1007/S00338-021-02064-3/TABLES/1

Moynihan, M.A., Amini, S., Oalmann, J., Chua, J.Q.I., Tanzil, J.T.I., Fan, T.Y., Miserez, A., Goodkin, N.F., 2022. Crystal orientation mapping and microindentation reveal anisotropy in Porites skeletons. Acta Biomater. 151, 446–456. 10.1016/J.ACTBIO.2022.08.012

Mwachireya, S.A., McClanahan, T.R., Cote, I.M., Hartwick, B.E., 2016. Increased Terrestrial Perturbations Modify Skeletal Properties and Mechanical Strength of Hard Corals. Environment and Natural Resources Research 6, 153. 10.5539/ENRR.V6N4P153

Nakamura, T., Van Woesik, R., 2001. Water-flow rates and passive diffusion partially explain differential survival of corals during the 1998 bleaching event. Mar. Ecol. Prog. Ser. 212, 301–304. 10.3354/MEPS212301

Omer, M., Carrasco-Pena, A., Orlovskaya, N., Collins, B.E., Yarmolenko, S.N., Sankar, J., Subhash, G., Gilliam, D.S., Fauth, J.E., 2020. Structural and mechanical properties of staghorn coral (Acropora cervicornis) CaCO3 aragonite skeletons, cleaned by chemical bleaching and biological processes. Advances in Applied Ceramics 119, 434–438. 10.1080/17436753.2020.1815500;CTYPE:STRING:JOURNAL

Palumbi, S.R., Barshis, D.J., Traylor-Knowles, N., Bay, R.A., 2014. Mechanisms of reef coral resistance to future climate change. Science (1979). 344, 895–898. 10.1126/science.1251336

Pasquini, L., Molinari, A., Fantazzini, P., Dauphen, Y., Cuif, J.P., Levy, O., Dubinsky, Z., Caroselli, E., Prada, F., Goffredo, S., Di Giosia, M., Reggi, M., Falini, G., 2015. Isotropic microscale mechanical properties of coral skeletons. J. R. Soc. Interface 12. 10.1098/rsif.2015.0168

Paz-García, D.A., Aldana-Moreno, A., Cabral-Tena, R.A., García-De-León, F.J., Hellberg, M.E., Balart, E.F., 2015. Morphological variation and different branch modularity across contrasting flow conditions in dominant Pocillopora reef-building corals. Oecologia 178, 207–218. 10.1007/s00442-014-3199-9

Perry, C.T., Alvarez-Filip, L., Graham, N.A.J., Mumby, P.J., Wilson, S.K., Kench, P.S., Manzello, D.P., Morgan, K.M., Slangen, A.B.A., Thomson 11, D.P., Januchowski-Hartley, F., Smithers, S.G., John’, S., Labrador, C., 2018. Loss of coral reef growth capacity to track future increases in sea level. Nature 558, 396–400. 10.1038/s41586-018-0194-z

Prada, F., Caroselli, E., Mengoli, S., Brizi, L., Fantazzini, P., Capaccioni, B., Pasquini, L., Fabricius, K.E., Dubinsky, Z., Falini, G., Goffredo, S., 2017. Ocean warming and acidification synergistically increase coral mortality. Sci. Rep. 7. 10.1038/srep40842

Richardson, L.E., Graham, N.A.J., Hoey, A.S., 2020. Coral species composition drives key ecosystem function on coral reefs. Proceedings of the Royal Society B: Biological Sciences 287, 20192214. 10.1098/RSPB.2019.2214

Roche, R.C., Abel, R.A., Johnson, K.G., Perry, C.T., 2010. Quantification of porosity in Acropora pulchra (Brook 1891) using X-ray micro-computed tomography techniques. J. Exp. Mar. Biol. Ecol. 396, 1–9. 10.1016/J.JEMBE.2010.10.006

Schueth, J.D., Frank, T.D., 2008. Reef foraminifera as bioindicators of coral reef health: Low Isles reef, northern Great Barrier Reef, Australia. The Journal of Foraminiferal Research 38, 11–22. 10.2113/gsjfr.38.1.11

Scucchia, F., Zaslansky, P., Boote, C., Doheny, A., Mass, T., Camp, E.F., 2023. The role and risks of selective adaptation in extreme coral habitats. Nature Communications 2023 14:1 14, 1–12. 10.1038/s41467-023-39651-7

Tambutté, E., Venn, A.A., Holcomb, M., Segonds, N., Techer, N., Zoccola, D., Allemand, D., Tambutté, S., 2015. Morphological plasticity of the coral skeleton under CO2-driven seawater acidification. Nature Communications 6. 10.1038/ncomms8368

Todd, P.A., 2008. Morphological plasticity in scleractinian corals. Biological Reviews 83, 315–337. 10.1111/J.1469-185X.2008.00045.X;PAGEGROUP:STRING:PUBLICATION

Waller, A., Byrne, M., Foo, S.A., 2025. Coexisting Mangrove-Coral Habitats: Trends in Seawater Chemistry and Implications for Corals. Oceanography and Marine Biology: An Annual Review 97–113. 10.1201/9781003589600-3

Wang, J., Zhou, B., Liu, X.S., Fields, A.J., Sanyal, A., Shi, X., Adams, M., Keaveny, T.M., Guo, X.E., 2015. Trabecular plates and rods determine elastic modulus and yield strength of human trabecular bone. Bone 72, 71–80. 10.1016/j.bone.2014.11.006

Xu, J., Zhang, Z., Huang, X., 2025. Progressive failure and fracture characteristics of coral reef limestone under triaxial compression. Eng. Fract. Mech. 318. 10.1016/j.engfracmech.2025.110948

Yates, K.K., Rogers, C.S., Herlan, J.J., Brooks, G.R., Smiley, N.A., Larson, R.A., 2014. Diverse coral communities in mangrove habitats suggest a novel refuge from climate change. Biogeosciences 11, 4321–4337. 10.5194/bg-11-4321-2014

