## Supplementary material for "Reduced skeletal density and mechanical resistance in mangrove-associated *Pocillopora* sp. corals"

**Table S1. Stiffness and regression quality for all 15 specimens**

| Site | Stiffness (N mm⁻¹) | $R^2$ | n (linear pts) | Peak force (N) | $R^2 < 0.98$† | Loading mode |
|---|---|---|---|---|---|---|
| ML | 728.7 | 0.973 | 32 | 334.2 | ✓ | Distance |
| ML | 565.3 | 0.837 | 18 | 237.2 | ✓ | Distance |
| ML | 1961.5 | 0.976 | 20 | 722.3 | ✓ | Force |
| ML | 652.0 | 0.987 | 50 | 517.5 | | Force |
| MC | 362.8 | 0.914 | 36 | ≥247.9 | ✓ | Distance |
| MC | 674.3 | 0.964 | 71 | ≥657.6 | ✓ | Distance |
| MC | 918.1 | 0.939 | 14 | 291.1 | ✓ | Distance |
| MC | 261.6 | 0.773 | 45 | 301.9 | ✓ | Distance |
| DR | 398.8 | 0.926 | 33 | ≥258.7 | ✓ | Distance |
| DR | 448.7 | 0.981 | 60 | ≥377.3 | | Force |
| DR | 656.5 | 0.926 | 49 | 474.3 | ✓ | Distance |
| PR | 2634.3 | 0.996 | 87 | ≥3331.2 | | Distance |
| PR | 1941.8 | 0.991 | 80 | 2016.0 | | Distance |
| PR | 1912.3 | 0.987 | 79 | ≥2113.0 | | Distance |
| PR | 381.5 | 0.873 | 68 | 495.9 | ✓ | Distance |

† $R^2$ values in bold indicate samples where the linear elastic region regression fell below $R^2 = 0.98$, suggesting the 20-70% force window may span a non-linear region. Stiffness = OLS regression slope on data within 20-70% of maximum force in the loading phase. Peak forces preceded by ≥ denote censored specimens (lower bound on true failure force). n = number of data points in the linear region. ML = Mangrove Lagoon, MC= Mangrove Channel, DR= Distal Reef, PR= Proximal Reef

**Table S2. Baseline (T1) values for all eleven CTAn morphometric parameters by site**

***(a) Volume, surface, and density parameters***

| Parameter | ML | MC | DR | PR |
|---|---|---|---|---|
| S/V ratio ($mm^{-1}$) | 7.52 ± 2.29 | 7.00 ± 0.85 | 4.23 ± 1.13 | 1.96 ± 0.67 |
| % Object volume | 7.26 ± 3.04 | 5.88 ± 1.35 | 10.96 ± 3.46 | 18.80 ± 4.71 |
| Object volume ($mm^3$) | 158.19 ± 69.50 | 172.20 ± 54.49 | 268.98 ± 73.80 | 258.78 ± 81.69 |
| Object surface ($mm^2$) | 1168.95 ± 482.78 | 1209.20 ± 399.81 | 1103.93 ± 284.17 | 475.07 ± 74.05 |
| Intersection surface ($mm^2$) | 16.73 ± 4.74 | 13.70 ± 3.68 | 21.20 ± 8.65 | 33.67 ± 5.99 |
| Object surface density | 0.494 ± 0.089 | 0.414 ± 0.115 | 0.443 ± 0.106 | 0.351 ± 0.062 |
| Structure linear density | 0.139 ± 0.040 | 0.121 ± 0.033 | 0.113 ± 0.033 | 0.103 ± 0.019 |

ML = Mangrove Lagoon, MC= Mangrove Channel, DR= Distal Reef, PR= Proximal Reef

***(b) Structural thickness and separation parameters***

| Parameter | ML | MC | DR | PR |
|---|---|---|---|---|
| Structure thickness (mm) | 0.507 ± 0.091 | 0.492 ± 0.053 | 1.018 ± 0.433 | 1.874 ± 0.516 |
| Structure separation (mm) | 5.523 ± 1.381 | 6.026 ± 0.587 | 4.992 ± 0.761 | 3.664 ± 0.413 |
| SD structure thickness (mm) | 0.223 ± 0.048 | 0.199 ± 0.010 | 0.441 ± 0.191 | 0.716 ± 0.167 |
| SD structure separation (mm) | 1.894 ± 0.484 | 1.940 ± 0.177 | 1.929 ± 0.210 | 1.058 ± 0.087 |

Values are mean ± SD. ML = Mangrove Lagoon, MC= Mangrove Channel, DR= Distal Reef, PR= Proximal Reef

**Table S3. Kruskal-Wallis test results for baseline (T1) differences among sites for all eleven CTAn morphometric parameters.**

| Parameter | H | df | p | Sig. |
|---|---|---|---|---|
| Object volume ($mm^3$) | 6.079 | 3 | 0.1078 | ns |
| % Object volume | 10.413 | 3 | 0.0154 | * |
| Object surface ($mm^2$) | 7.013 | 3 | 0.0715 | ns |
| Intersection surface ($mm^2$) | 8.779 | 3 | 0.0324 | * |
| S/V ratio ($mm^{-1}$) | 11.029 | 3 | 0.0116 | * |
| Object surface density | 4.225 | 3 | 0.2382 | ns |
| Structure thickness (mm) | 11.329 | 3 | 0.0101 | * |

| Parameter | H | df | p | Sig. |
|---|---|---|---|---|
| Structure separation (mm) | 8.929 | 3 | 0.0302 | * |
| Structure linear density | 1.717 | 3 | 0.6332 | ns |
| SD structure thickness (mm) | 10.825 | 3 | 0.0127 | * |
| SD structure separation (mm) | 8.475 | 3 | 0.0372 | * |

H = Kruskal-Wallis chi-squared statistic; df = degrees of freedom. Significance: * $p < 0.05$, ns = not significant.

**Table S4. Dunn's post-hoc pairwise comparisons (Bonferroni-corrected) for the seven baseline parameters with significant Kruskal-Wallis results.**

***S/V ratio (mm$^{-1}$)***

| Comparison | z | p (Bonferroni) | Sig. |
|---|---|---|---|
| ML vs MC | 0.079 | 1.0000 | ns |
| ML vs DR | 1.513 | 0.7822 | ns |
| ML vs PR | 2.846 | 0.0266 | * |
| MC vs DR | 1.439 | 0.9001 | ns |
| MC vs PR | 2.767 | 0.0339 | * |
| DR vs PR | 1.122 | 1.0000 | ns |

ML = Mangrove Lagoon, MC= Mangrove Channel, DR= Distal Reef, PR= Proximal Reef

***% Object volume***

| Comparison | z | p (Bonferroni) | Sig. |
|---|---|---|---|
| ML vs MC | 0.553 | 1.0000 | ns |
| ML vs DR | -0.952 | 1.0000 | ns |
| ML vs PR | -2.451 | 0.0855 | ns |
| MC vs DR | -1.464 | 0.8594 | ns |
| MC vs PR | -3.004 | 0.0160 | * |
| DR vs PR | -1.317 | 1.0000 | ns |

ML = Mangrove Lagoon, MC= Mangrove Channel, DR= Distal Reef, PR= Proximal Reef

***Structure thickness (mm)***

| Comparison | z | p (Bonferroni) | Sig. |
|---|---|---|---|
| ML vs MC | 0.316 | 1.0000 | ns |
| ML vs DR | -1.561 | 0.7105 | ns |
| ML vs PR | -2.609 | 0.0545 | ns |
| MC vs DR | -1.854 | 0.3823 | ns |
| MC vs PR | -2.925 | 0.0207 | * |

| Comparison | z | p (Bonferroni) | Sig. |
|---|---|---|---|
| DR vs PR | -0.854 | 1.0000 | ns |

ML = Mangrove Lagoon, MC= Mangrove Channel, DR= Distal Reef, PR= Proximal Reef

***Structure separation (mm)***

| Comparison | z | p (Bonferroni) | Sig. |
|---|---|---|---|
| ML vs MC | -0.949 | 1.0000 | ns |
| ML vs DR | 0.195 | 1.0000 | ns |
| ML vs PR | 1.976 | 0.2886 | ns |
| MC vs DR | 1.073 | 1.0000 | ns |
| MC vs PR | 2.925 | 0.0207 | * |
| DR vs PR | 1.635 | 0.6128 | ns |

ML = Mangrove Lagoon, MC= Mangrove Channel, DR= Distal Reef, PR= Proximal Reef

***Intersection surface (mm²)***

| Comparison | z | p (Bonferroni) | Sig. |
|---|---|---|---|
| ML vs MC | 0.474 | 1.0000 | ns |
| ML vs DR | -0.683 | 1.0000 | ns |
| ML vs PR | -2.293 | 0.1312 | ns |
| MC vs DR | -1.122 | 1.0000 | ns |
| MC vs PR | -2.767 | 0.0339 | * |
| DR vs PR | -1.439 | 0.9001 | ns |

ML = Mangrove Lagoon, MC= Mangrove Channel, DR= Distal Reef, PR= Proximal Reef

***SD structure thickness (mm)***

| Comparison | z | p (Bonferroni) | Sig. |
|---|---|---|---|
| ML vs MC | 0.395 | 1.0000 | ns |
| ML vs DR | -1.391 | 0.9860 | ns |
| ML vs PR | -2.530 | 0.0685 | ns |
| MC vs DR | -1.757 | 0.4739 | ns |
| MC vs PR | -2.925 | 0.0207 | * |
| DR vs PR | -0.952 | 1.0000 | ns |

ML = Mangrove Lagoon, MC= Mangrove Channel, DR= Distal Reef, PR= Proximal Reef

***SD structure separation (mm)***

| Comparison | z | p (Bonferroni) | Sig. |
|---|---|---|---|
| ML vs MC | -0.474 | 1.0000 | ns |
| ML vs DR | -0.220 | 1.0000 | ns |

| Comparison | z | p (Bonferroni) | Sig. |
|---|---|---|---|
| ML vs PR | 2.135 | 0.1968 | ns |
| MC vs DR | 0.220 | 1.0000 | ns |
| MC vs PR | 2.609 | 0.0545 | ns |
| DR vs PR | 2.196 | 0.1686 | ns |

z = Dunn's test statistic; p (Bonferroni) = two-tailed p-value adjusted for six pairwise comparisons per parameter. Significance: * $p < 0.05$, ns = not significant. ML = Mangrove Lagoon, MC= Mangrove Channel, DR= Distal Reef, PR= Proximal Reef

**Table S5. Correlation of stiffness with all eleven baseline (T1) morphometric parameters.**

| Morphometric parameter | Pearson r | Pearson p | Pearson p (adj.) | Spearman ρ | Spearman p | Spearman p (adj.) | Sig. |
|---|---|---|---|---|---|---|---|
| SD structure separation (mm) | -0.688 | 0.005 | 0.019 | -0.571 | 0.029 | 0.195 | * |
| Object surface area ($mm^2$) | -0.681 | 0.005 | 0.019 | -0.489 | 0.067 | 0.195 | * |
| Structure thickness (mm) | 0.680 | 0.005 | 0.019 | 0.482 | 0.071 | 0.195 | * |
| % Object volume | 0.645 | 0.009 | 0.022 | 0.457 | 0.089 | 0.195 | * |
| Intersection surface area ($mm^2$) | 0.642 | 0.010 | 0.022 | 0.411 | 0.130 | 0.204 | * |
| Structure separation (mm) | -0.607 | 0.016 | 0.028 | -0.425 | 0.116 | 0.204 | * |
| SD structure thickness (mm) | 0.594 | 0.020 | 0.028 | 0.321 | 0.242 | 0.333 | * |
| S/V ratio ($mm^{-1}$) | -0.592 | 0.020 | 0.028 | -0.496 | 0.062 | 0.195 | * |
| Object surface density ($mm^2/mm^3$) | -0.320 | 0.244 | 0.298 | -0.204 | 0.466 | 0.569 | ns |
| Object volume ($mm^3$) | 0.185 | 0.509 | 0.560 | 0.111 | 0.695 | 0.765 | ns |
| Structure linear density ($mm^{-1}$) | -0.102 | 0.718 | 0.718 | -0.068 | 0.812 | 0.812 | ns |

Parameters are ordered by the magnitude of the Pearson correlation coefficient (r). p (adj.) = p-value adjusted for multiple comparisons across all eleven parameters using the Benjamini-Hochberg procedure. Sig. indicates significance of the p-value: * $p < 0.05$; ns = not significant. n = 15 for all correlations.
